# A Double-Blind Pilot Dosing Study of Low Field Magnetic Stimulation (LFMS) for Treatment-Resistant Depression (TRD)

**DOI:** 10.1101/465013

**Authors:** Marc J. Dubin, Irena Ilieva, Zhi-De Deng, Jeena Thomas, Ashly Cochran, Kamilla Kravets, Benjamin D. Brody, Paul J. Christos, James H. Kocsis, Conor Liston, Faith M. Gunning

## Abstract

Low Field Magnetic Stimulation is a potentially rapid-acting treatment for depression with mood-enhancing effects in as little as one 20-minute session. The most convincing data for LFMS has come from treating bipolar depression. We examined whether LFMS also has rapid mood-enhancing effects in treatment-resistant major depressive disorder, and whether these effects are dose-dependent. We hypothesized that a single 20-min session of LFMS would reduce depressive symptom severity and that the magnitude of this change would be greater after three 20-min sessions than after a single 20-min session. In a double-blind randomized controlled trial, 30 participants (age 21–65) with treatment-resistant depression were randomized to three 20-minute active or sham LFMS treatments with 48 hours between treatments. Response was assessed immediately following LFMS treatment using the 6-item Hamilton Depression Rating Scale (HAMD-6), the Positive and Negative Affect Scale (PANAS) and the Visual Analog Scale. Following the third session of LFMS, the effect of LFMS on VAS and HAMD-6 was superior to sham (*F*(1, 24) = 7.45, *p* = 0.03, Holm-Bonferroni corrected; *F*(1,22) = 6.92, *p* = 0.03, Holm-Bonferroni corrected, respectively). There were no differences between sham and LFMS following the initial or second session with the effect not becoming significant until after the third session. Three 20-minute LFMS sessions were required for active LFMS to have a mood-enhancing effect for individuals with treatment-resistant depression. As this effect may be transient, future work should address dosing schedules of longer treatment course as well as biomarker-based targeting of LFMS to optimize patient selection and treatment outcomes.

## I. Introduction

Treatment-resistant depression (TRD) is a significant clinical problem with approximately two-thirds of patients with major depressive disorder (MDD) not achieving remission after the first trial of antidepressant medications and one-third of patients remaining symptomatic following multiple sequential trials of antidepressant medication [1]. Other patients are unable to tolerate antidepressant medications [2, 3]. Although treatments such as electroconvulsive therapy (ECT) are highly efficacious [4–6], tolerability of ECT can be low due to potential cognitive side effects [7]. Newer magnetic stimulation treatments such as repetitive transcranial magnetic stimulation (rTMS) have shown clinical efficacy, reproduced in meta-analyses [8–10], and have a better side effect profile than ECT and many medications. However, the standard rTMS treatment protocol requires daily 40-minute sessions, continuously for 4–6 weeks, and therefore is inconvenient and costly for patients.

Low field magnetic stimulation may offer an alternative to patients who do not remit with traditional pharmacologic approaches. LFMS employs the unique magnetic field waveform used in echo-planar magnetic resonance spectroscopic imaging to deliver a low intensity, time-varying electric field in the brain (*E* ≤ 1V/m, 1 kHz). Unlike TMS, LFMS does not evoke neuronal action potentials, but has been shown to modulate metabolism in broad regions of human cerebral cortex [11], and to modulate spontaneous neuronal oscillations in rodent slice preparations [12, 13]. LFMS also had significant antidepressant-like behavioral effects in a rodent model, reducing immobility in a forced swim test protocol [14]. However, mood enhancement has been equivocal in humans with MDD, with recent single and multi-site randomized controlled trials missing targets on their primary outcome variables, while showing some positive results on select secondary outcomes [15, 16]. This is in contrast to somewhat more consistent positive mood-enhancing results for bipolar depression [16, 17]. LFMS may have safety advantages over TMS that could broaden its clinical applicability. Because it is subthreshold, it does not require periodic monitoring of motor threshold. Therefore, it is likely to be better tolerated than TMS, carry a lower seizure risk, and thus could potentially be administered in an unsupervised setting.

The main objectives of this study were to examine the mood effects of LFMS on patients with TRD and to explore dosing in terms of number of treatment sessions. To this end, we undertook a double-blind randomized controlled trial of 3 sessions of LFMS for TRD. We hypothesized that relative to participants randomized to the sham condition, measures of mood would be greater in the active LFMS group immediately after session 3. In an exploratory analysis, we investigated whether a difference in measures of mood emerged earlier, after the first or second treatment session.

## II. Methods

This was a single-site, double-blind, randomized, sham-controlled phase II study of the efficacy of LFMS on mood symptoms in TRD. Eligible participants were randomly assigned to double-blind treatment with three 20-minute sessions of either active or sham LFMS. This trial was conducted according to the U.S. Food and Drug Administration guidelines (https://clinicaltrials.gov/ct2/show/NCT01944644) and the Declaration of Helsinki. Written informed consent was obtained from all participants before protocol-specified procedures were carried out. The participants were drawn from an outpatient sample of patients with current MDD. All aspects of our experimental protocol were approved by the Institutional Review Board of Weill Cornell Medical College and conducted in accordance with institutional guidelines.

## A. Participants

The Principal Investigator (MJD) screened participants for eligibility criteria (below) as outlined in the Study Flowchart (Figure 1). In cases in which the participant was unable to answer all relevant questions regarding their psychiatric history, additional history was obtained from current and past psychiatric treaters. A HIPAA release was obtained from the participant when this additional history was required. Treatment-resistance was defined as the failure to respond to one or more trials of an adequate dose of an antidepressant for at least 8 weeks and was documented using the MGH Antidepressant Treatment Response Questionnaire (MGH ATRQ) [18–20]. In addition, if currently on psychotropic medication, participants must have been taking the current doses of all medications in their current regimen for the past four weeks [21, 22]. Thirty participants were recruited who met the following inclusion and exclusion criteria.

**Figure 1:**
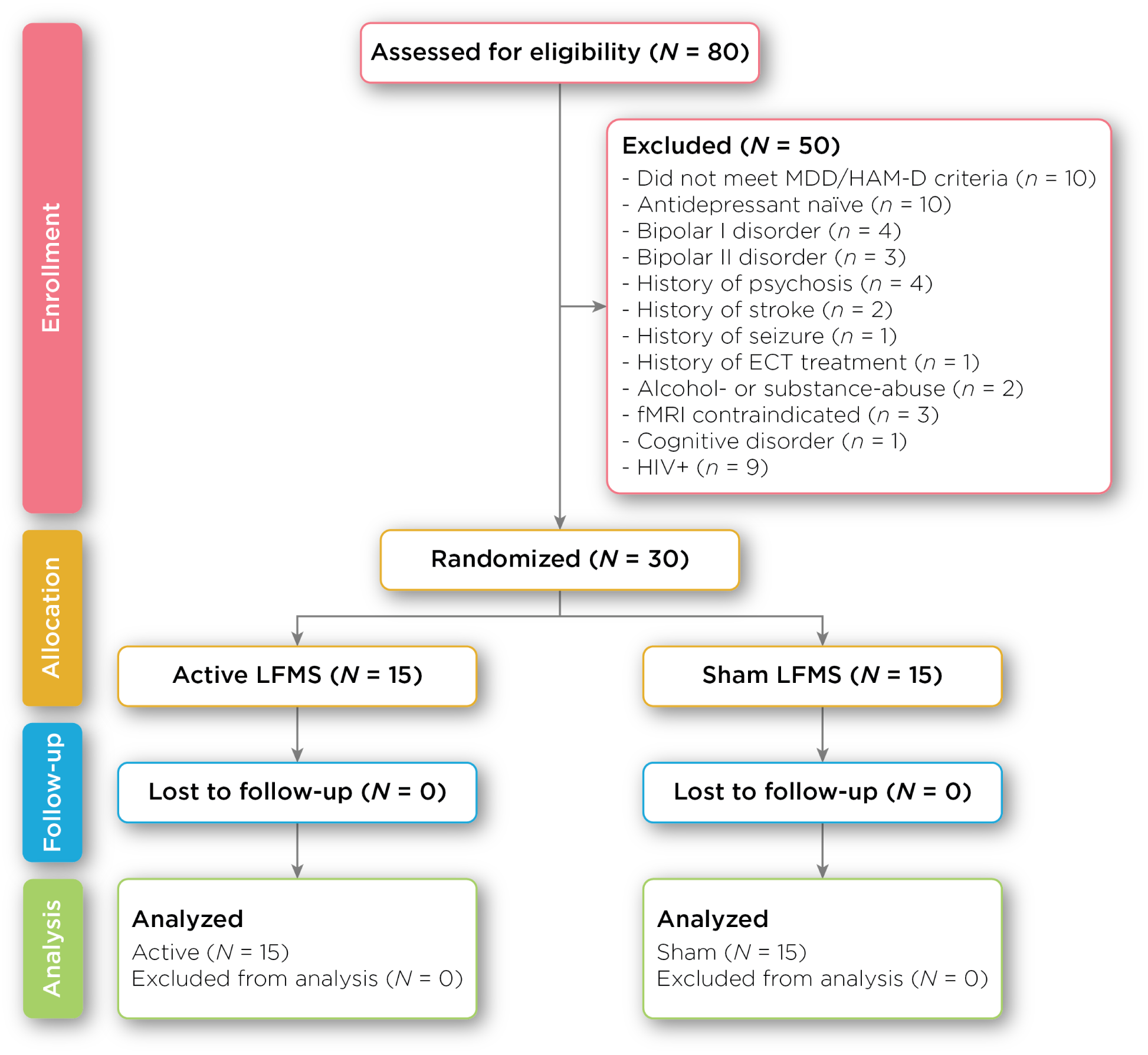
CONSORT Diagram of the LFMS protocol, showing number of participants from enrollment phase through analysis.

**Figure 2:**
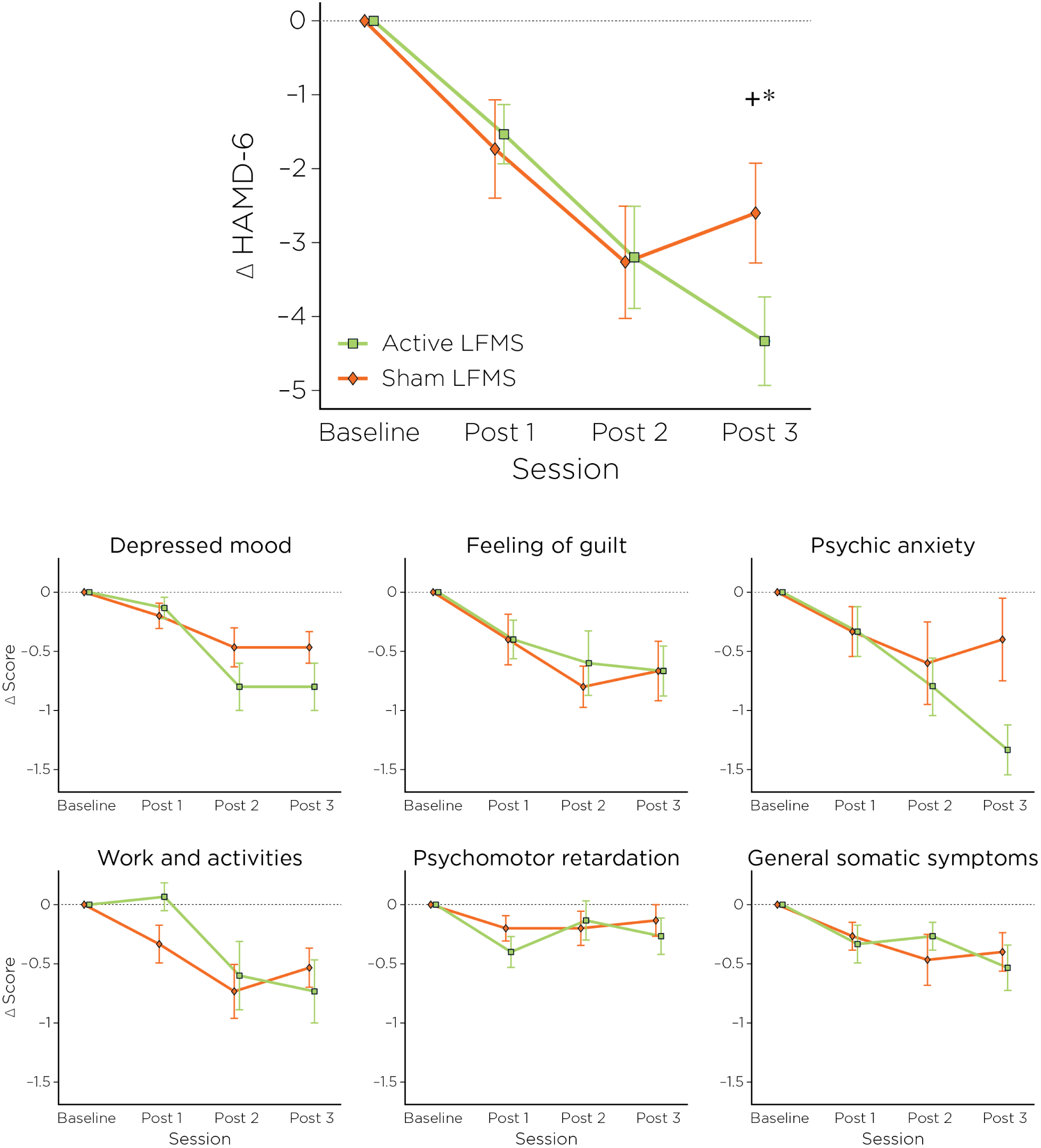
Effect of active vs. sham LFMS on symptoms of depression, measured by HAMD-6. The top graph shows the difference from baseline after active LFMS (green line) compared to sham LFMS (orange). The difference increases with subsequent sessions and is statistically significant after the final treatment. (^+^ uncorrected; ^∗^ Holm–Bonferroni corrected). Bottom panels show the effect of LFMS on each of the 6 item scores that comprise the HAMD-6. The largest effect of active LFMS is on the Depressed mood and Psychic anxiety items.

**Figure 3:**
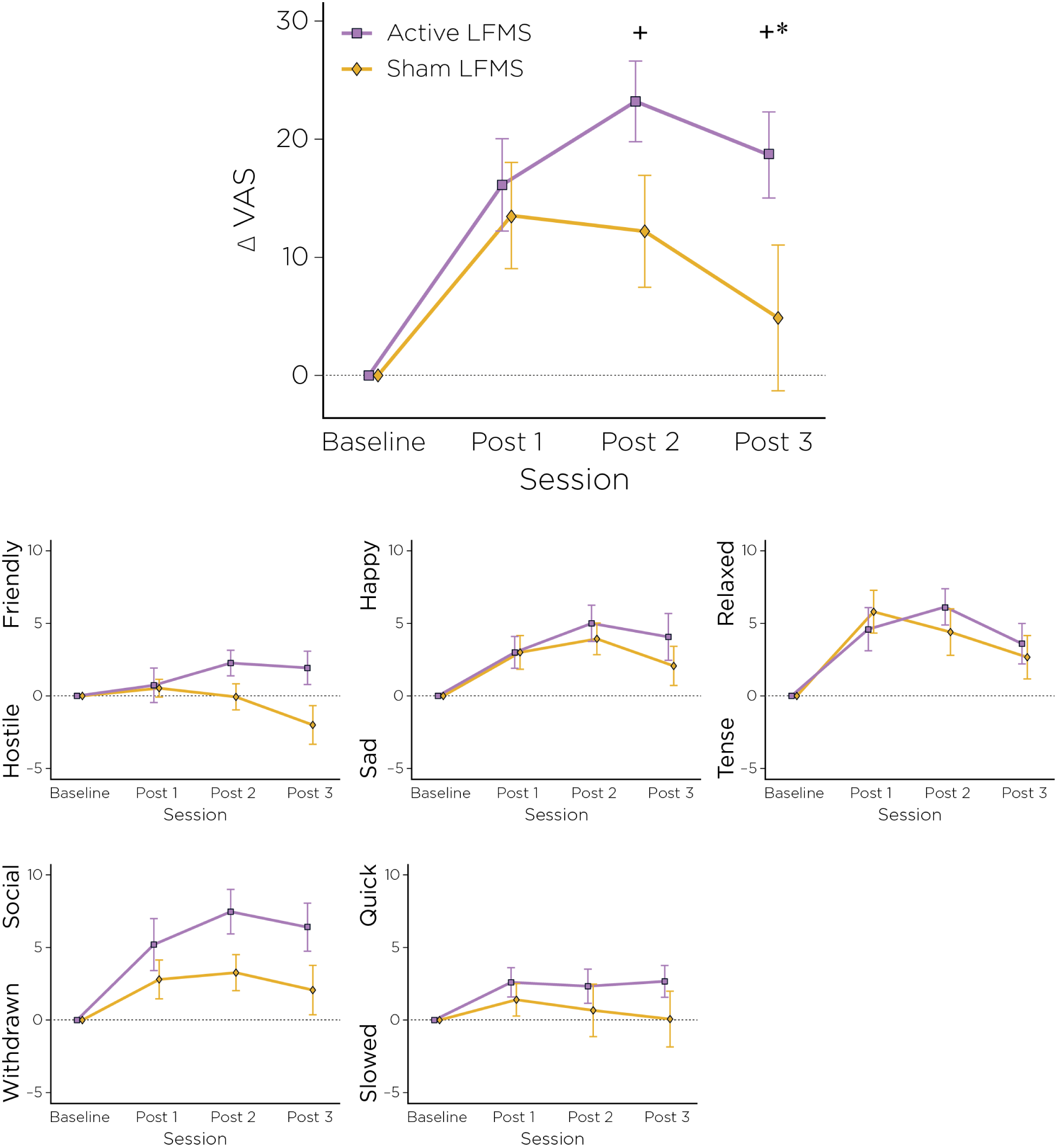
Effect of Active LFMS vs. Sham on mood symptoms, measured by VAS. The top graph shows the difference from baseline after active LFMS (purple line) compared to sham LFMS (yellow). The difference increases with subsequent sessions and is statistically significant after the final treatment. (^+^ uncorrected ^∗^ Holm–Bonferroni corrected). Bottom panels show effect of LFMS on each of the 5 item scores that comprise the VAS. The largest effect of active LFMS is on the Hostile-Friendly and Withdrawn–Social items.

### 1) Inclusion Criteria

A participant was eligible for participation in the study if all of the following criteria were met:

a. Able to understand and read English and give written informed consent prior to the protocol required procedures.
b. Men and women, ages 18 to 65 inclusive with a diagnosis of major depressive episode as defined by DSM-IV-TR criteria and diagnosed by a board-certified psychiatrist (MJD) using the Mini International Neuropsychiatric Interview (MINI) Version 5.
c. Inadequate response to 1 or more adequate antidepressant treatments in the current depressive episode.
d. 17-item Hamilton Rating Scale for Depression (HAMD-17) score ≥ 18 administered by a research assistant certified in the administration of the HAMD.
e. Body Mass Index (BMI) of approximately 18–40 kg/m^2^.
f. Women of childbearing potential (WOCBP) were required to be using an adequate method of contraception to avoid pregnancy throughout the study and must have had a negative urine pregnancy test within 72 hours prior to the start of LFMS.

#### 2) Exclusion Criteria

A participant was ineligible for participation in the study if any of the following criteria were met:

a. WOCBP who were unwilling or unable to use an acceptable method to avoid pregnancy for the entire study period.
b. Women who were pregnant or breastfeeding.
c. Participants with other DSM-IV-TR Axis I disorders other than Generalized Anxiety Disorder (GAD: 300.02), Social Anxiety Disorder (300.23), or Specific Phobia (300.29). Participants with co-morbid GAD, Social Anxiety Disorder, or Specific Phobia are ineligible if the co-morbid condition was clinically unstable, required treatment, or had been the primary focus of treatment within the 6-month period prior to screening.
d. Delirium, dementia, or other cognitive disorder.
e. Schizophrenia or other psychotic disorder, based on the MINI.
f. Patients with a clinically significant Axis II diagnosis of borderline, antisocial, paranoid, schizoid, schizotypal or histrionic personality disorder.
g. Patients experiencing hallucinations, delusions, or any psychotic symptomatology in the current or any previous depressive episode.
h. Patients who met DSM-IV-TR criteria for any significant substance use disorder within the six months prior to screening.
i. Patients who received new-onset psychotherapy and/or somatic therapy (light therapy, transcranial magnetic stimulation) within 6 weeks of screening, or at any time during participation in the trial.
j. Patients who, in the opinion of the Investigator, were actively suicidal and at significant risk for suicide.
k. Patients who had participated in any clinical trial with an investigational drug or device within the month prior to screening.
l. Patients who had received ECT in the past 20 years or Vagus Nerve/Deep Brain Stimulation during their lifetime.
m. Patients who had medical illness including, cardiovascular, hepatic, renal, respiratory, endocrine, neurological, or hematological disease.
n. Participants with evidence or history of significant neurological disorder, including head trauma with loss of consciousness, history of stroke, Parkinson’s disease, epilepsy disorder, conditions that lower seizure threshold, seizures of any etiology (including substance or drug withdrawal), who were taking medications to control seizures, or who had increased risk of seizures as evidenced by history of EEG with epileptiform activity (with the exception of juvenile febrile seizures).
o. Patients with thyroid pathology (unless condition had been stabilized with medications for at least the three months prior to screening).
p. Patients who had begun any medications in the two weeks prior to screening.
q. Monoamine oxidase inhibitors (e.g., Nardil, phenelzine, Parnate, tranylcypromine, Marplan, isocarboxazide) treatment within the 2 weeks prior to enrollment.
r. Patients who had a history of antidepressant-induced hypomania or dysphoria.
s. Participants who had metal implants (Defined by the NY Presbyterian Hospital MRI Checklist).

### B. Study Design

All participants in the intent-to-treat group were randomized with a 1: 1 ratio to LFMS or sham. All LFMS sessions were 20 min in duration. (1) Each participant in the active LFMS group received 3 days of LFMS treatment, with each treatment session consisting of 20 min of active LFMS. (2) The sham group received 3 days of sham LFMS treatment, with each treatment session consisting of 20 min of sham LFMS. For both active and sham groups, successive treatments were separated by approximately 48 hours.

Mood-enhancing response was evaluated by assessing changes in three primary outcome measures immediately prior to and immediately after each LFMS or sham session. The three measures were: the observer-rated 6-item Hamilton Depression Rating Scale (HAMD-6) [23, 24], the participant-rated Positive and Negative Affect Scale (PANAS) [25], and the participant-rated Visual-Analog Mood Scale (VAS), assessing sad-happy, withdrawn-social, tense-relaxed, slow-quick, and hostile-friendly. The VAS score was defined as the average of these five components [26–28]. The HAMD-6 ratings were conducted by two research assistants trained by certified HAMD raters (MJD and JK).

If the HAMD-6 declined by 25% or greater, either between successive treatments or from before to after a given treatment, the participant was terminated from the study. The Principal Investigator evaluated the participant for acute risk of suicide. If the participant was determined to be at acute risk for suicide, the participant was brought to the psychiatric emergency room at Weill Cornell for further evaluation. Follow-up care with the participant’s outside psychiatrist was arranged by the Principal Investigator.

### C. LFMS Protocol

LFMS followed a previously published protocol for the treatment of a Major Depressive Episode [17]. LFMS treatments were delivered with a prototype LFMS device manufactured by Tal Medical (Boston, MA). LFMS sessions consisted of a head coil and pulse sequence as used in proton echo-planar magnetic resonance spectroscopic imaging and were 20 min in duration. LFMS exposes participants to magnetic fields of the same magnitude and frequency used in clinical MR-Spectroscopic imaging of the brain. Sham LFMS consisted of a three-dimensional spoiled gradient echo sequence of the same duration as active LFMS and which provided auditory stimulation indistinguishable from active treatment.

At the beginning of a treatment session, the participant lay on a flat, padded table and positioned his head within the open bore of the LFMS device such that his superior orbital ridge aligned with the opening of the bore. The device was preprogrammed to deliver active or sham treatment so that the participant, operator, and all investigators were blinded to active treatment vs. sham. Immediately before and after each treatment session, the HAMD-6, PANAS, and VAS were administered and the participant was monitored for any adverse events.

### D. Adverse Events

Observed and spontaneously-reported adverse events were recorded at each visit. Spontaneously-reported adverse events were classified as mild, moderate or severe. Participants were allowed to contact the principal investigator or a member of the study team at any time between visits concerning adverse events or worsening of symptoms.

### E. Statistical Analysis Plan

Descriptive statistics (including mean, standard deviation, frequency, and percent) for demographic and baseline clinical severity factors were calculated to characterize the active LFMS and sham groups (both for per-protocol and intent-to-treat). The two-sample *t*-test or *χ*^2^-test were used, as appropriate, to compare demographic and baseline clinical severity factors between the active LFMS and sham groups. Analysis of covariance (ANCOVA) was used to compare mood ratings (i.e., HAMD-6, PANAS-Positive, PANAS-Negative, and VAS; 4 separate ANCOVA models) at the final session between the active and sham groups, controlling for age, gender, and baseline mood score. In cases of missing data, the last-observation-carried-forward (LOCF) method was used in the ANCOVA analyses. All *p*-values are two-sided with statistical significance evaluated at the 0.05 alpha level. Both uncorrected *p*-values and Holm–Bonferroni corrected *p*-values are presented for the ANCOVA models. For the intent-to-treat analysis, a sample size of 30 randomly assigned eligible patients in each group was calculated to have 80% power to detect a difference in means of 5.5 on the post-treatment HAMD-6 (between the active LFMS group and the sham group), assuming a common standard deviation of 7.0, using a two-group *t*-test with a two-sided alpha level of 5%. This calculation allowed for a 10% dropout rate. All analyses were performed in SPSS Version 24.0 (IBM Corp. Released 2016. IBM SPSS Statistics for Windows, Version 24.0, Armonk, NY: IBM Corp.).

## III. Results

### A. Demographics and Sample Characteristics

Table 1 shows the demographic characteristics and baseline clinical severity of the sample. Active LFMS and sham groups did not differ significantly in age, gender, race or baseline clinical severity measured by any of the 3 primary outcome variables.

**Table 1:** Demographic information and baseline clinical severity

|  | Active (*n* = 15) |  | Sham (*n* = 15) |  | *p* |
| --- | --- | --- | --- | --- | --- |
|  | M | SD | M | SD |  |
| **Demographics** |  |  |  |  |  |
| Age | 48.4 | 14.8 | 39.9 | 12.4 | 0.10 |
| Gender (% female) | 60 |  | 60 |  |  |
| Race (%) |  |  |  |  |  |
| White | 73 |  | 53 |  |  |
| African American | 13 |  | 13 |  |  |
| Asian | 0 |  | 7 |  |  |
| Hispanic | 7 |  | 20 |  |  |
| Other | 7 |  | 7 |  |  |
| **Baseline Clinical Severity** |  |  |  |  |  |
| HAMD-17 | 21.9 | 4.0 | 23.5 | 3.3 | 0.23 |
| HAMD-6 | 12.7 | 1.6 | 11.9 | 1.7 | 0.19 |
| VAS | 50.7 | 21.3 | 38.4 | 17.2 | 0.09 |
| PANAS–Negative | 24.5 | 7.7 | 31.1 | 9.0 | 0.04<sup>†</sup> |

P-values are for Student's $t$ -tests
Active and sham groups did not differ on any demographic variables. Additionally, baseline clinical severity did not differ significantly between active and sham groups with the exception of the sham group having higher baseline PANAS–Negative scores
<sup>†</sup> Significant difference between LFMS and sham (uncorrected)

### B. LFMS Effects on Mood

Table 2 shows the means and standard deviations of mood ratings (HAMD-6, PANAS-Positive, PANAS-Negative, and VAS) in the active-LFMS and sham groups at baseline and upon completion of the third and final treatment session. To examine whether mood ratings differed between active and sham LFMS at the conclusion of treatment, we conducted three ANCOVAs (analyses of covariance), with intervention (active, sham) as a between-participants factor and age, sex and baseline mood rating as covariates. The three dependent variables were scores for the primary outcome variables (HAMD-6, PANAS-Negative, and VAS) post-intervention. The effect of active LFMS on VAS and HAMD-6 emerged superior to sham (*F*(1,24) = 7.45, *p* = 0.03, Holm–Bonferroni corrected; *F*(1, 22) = 6.92, *p* = 0.03, Holm–Bonferroni corrected, respectively).

**Table 2:**
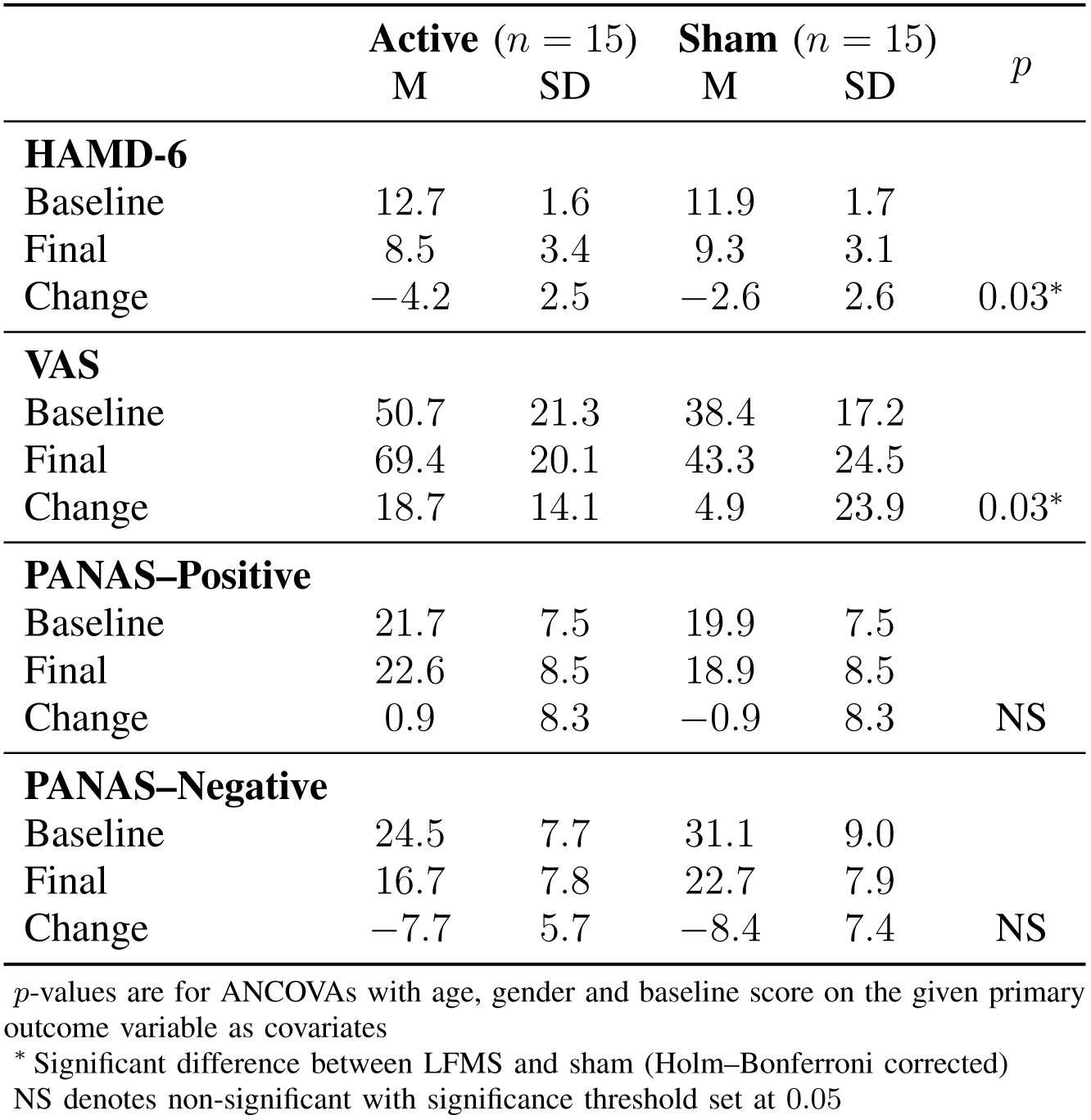
Effect of three 20 min sessions of active or sham LFMS on different primary outcome variables of mood

Additionally, to address dosing, we asked how early in the course of treatment significant differences emerge between the mood effects of active LFMS and sham. We conducted ANCOVAs with intervention (active, sham) as a between-participants factor and age, sex and baseline mood rating as covariates. The dependent variables were mood scores (HAMD-6, PANAS-Negative, and VAS) during the two mid-intervention (following the second LFMS session) assessment points. Among these tests, the mid-treatment differences between active LFMS and sham did not reach significance on HAMD-6, PANAS-Negative, or VAS after correcting for multiple comparisons.

### C. Effectiveness of Sham

For the 24 participants who completed a questionnaire about whether they believed they were in the active LFMS or sham condition, participants were not more likely than chance to correctly identify the intervention condition they were blind to (*χ*^2^ = 0.001, *p* = 0.97).

### D. Safety and Tolerability of LFMS

LFMS was tolerable. One participants dropped out after the first treatment because of serious clinical events, but these were deemed to be unrelated to the treatment itself. We observed no treatment-emergent suicidality. One subject who received sham was hospitalized 2 weeks after completion of the study for worsening depression and suicidal ideation.

## IV. Discussion

The principal finding of this study is that in a double-blinded sham-controlled study, 2 of 3 primary outcome variables of mood symptoms, namely the HAMD-6 and VAS, showed greater improvement after three treatments in the active LFMS group than in sham LFMS. This is the first study to demonstrate mood-enhancing effects of LFMS in treatment-resistant unipolar depression. The fact that 3 treatment sessions were required for a significant mood-enhancing effect to emerge between active and sham LFMS for TRD suggests that unipolar TRD may require more treatments than bipolar depression for a similar response. This is in keeping with bipolar depression responding faster than unipolar depression to ECT [29, 30].

Mood enhancements were unlikely to be attributable to sham because of the high quality of the sham—participants from active and sham groups guessed that they received active treatment at chance. Additionally, the mood-enhancing effect of the sham waned with subsequent treatments, an effect observed in other double-blinded, sham-controlled trials of brain stimulation treatments, including TMS [31].

The main limitation of our study is the lack of follow-up mood assessments and the short duration between treatment completion and participant assessments. This leaves open the possibility that the mood-enhancing effects reported here are highly transient, perhaps waning within hours. However, the mood enhancing effects observed here for TRD warrant further investigation that includes follow-up assessments to examine durability. Further, an enhanced dosing study of LFMS for TRD, to elucidate the number of sessions required to demonstrate durable mood improvements is required. If LFMS were shown to have more sustained effects using such a protocol, it would be of significant clinical value as this treatment could be delivered with reduced direct clinical supervision.

In addition to the limited mood assessment window, the use of the HAMD-6, although consistent with previous studies of LFMS, was a limitation of this study because of the time frame it refers to. Although we changed the time frame from 7 days to 48 hrs, the HAMD has still been validated only for the 7-day time period and this may have been a less appropriate assessment than the VAS or PANAS to capture transient effects on mood.

In summary, LFMS may be a promising, well-tolerated treatment for TRD with a potentially rapid onset of action. In addition to optimizing the dosing protocol and testing the durability of the mood improvments, we need to better characterize the subgroup of patients who respond to LFMS. Using functional neuroimaging to understand the circuit abnormalities that are predictive of treatment response has shown great promise for TMS [32–38]. Mapping these abnormalities in LFMS responders could eventually help match the treatment to depressed patients most likely to benefit and will guide advancements in LFMS coil design to optimize treatment [39, 40].

## Acknowledgments

This work was supported by a National Institutes of Health Weill Cornell Clinical and Translational Science Center pilot grant to MD and PL (UL1 TR002384) and additional grant funding from TAL Medical, Inc. CL was supported by grants from the National Institutes of Health (K99 MH097822) and the DeWitt Wallace Reader’s Digest Foundation at Weill Cornell. FG was supported by a grant from the National Institutes of Health (R01 MH097735). JK, BB, and AC were supported by a grant from the Pritzker Neuropsychiatric Disorders Research Consortium. ZD and JT are supported by the NIMH Intramural Research Program.

## References

[1] A. J. Rush, M. H. Trivedi, S. R. Wisniewski, A. A. Nierenberg, J. W. Stewart, D. Warden, G. Niederehe, M. E. Thase, P. W. Lavori, B. D. Lebowitz, P. J. McGrath, J. F. Rosenbaum, H. A. Sackeim, D. J. Kupfer, J. Luther, and M. Fava, “Acute and longer-term outcomes in depressed outpatients requiring one or several treatment steps: a STAR^∗^D report.” Am J Psychiatry, vol. 163, no. 1, pp. 1905–1917, 2006.

[2] M. Fava, “Management of nonresponse and intolerance: switching strategies,” J Clin Psychiatry, vol. 61, Suppl 2, pp. 10–12, 2000.

[3] G. M. Murphy, Jr., C. Kremer, H. E. Rodrigues, and A. F. Schatzberg, “Pharmacogenetics of antidepressant medication intolerance,” Am J Psychiatry, vol. 160, no. 10, pp. 1830–1835, 2003.

[4] UK ECT Review Group, “Efficacy and safety of electroconvulsive therapy in depressive disorders: a systematic review and meta-analysis,” Lancet, vol. 361, no. 9360, pp. 799–808, 2003.

[5] W. T. Heijnen, T. K. Birkenhäger, A. I. Wierdsma, and W. W. van den Broek, “Antidepressant pharmacotherapy failure and response to subsequent electroconvulsive therapy: a meta-analysis,” J Clin Psychopharmacol, vol. 30, no. 5, pp. 616–619, 2010.

[6] L. van Diermen, S. van den Ameele, A. M. Kamperman, B. C. G. Sabbe, T. Vermeulen, D. Schrijvers, and T. K. Birkenhager, “Prediction of electroconvulsive therapy response and remission in major depression: meta-analysis,” Br J Psychiatry, vol. 212, no. 2, pp. 71–80, 2018.

[7] M. Semkovska and D. M. McLoughlin, “Objective cognitive performance associated with electroconvulsive therapy for depression: a systematic review and meta-analysis,” Biol Psychiatry, vol. 68, no. 6, pp. 568–577, 2010.

[8] M. T. Berlim, F. van den Eynde, S. Tovar-Perdomo, and Z. J. Daskalakis, “Response, remission and drop-out rates following high-frequency repetitive transcranial magnetic stimulation (rTMS) for treating major depression: a systematic review and meta-analysis of randomized, double-blind and sham-controlled trials,” Psycho Med, vol. 44, no. 2, pp. 225–239, 2014.

[9] I. P. Ilieva, G. S. Alexopoulos, M. J. Dubin, S. S. Morimoto, L. W. Victoria, and F. M. Gunning, “Age-related repetitive transcranial magnetic stimulation effects on executive function in depression: a systematic review,” Am J Geriatr Psychiatry, vol. 26, no. 3, pp. 334–346, 2018.

[10] J. Mutz, D. R. Edgcumbe, A. R. Brunoni, and C. H. Y. Fu, “Efficacy and acceptability of non-invasive brain stimulation for the treatment of adult unipolar and bipolar depression: A systematic review and meta-analysis of randomised sham-controlled trials,” Neurosci Biobehav Rev, vol. 92, pp. 291–303, 2018.

[11] N. D. Volkow, D. Tomasi, G. J. Wang, J. S. Fowler, F. Telang, R. Wang, D. Alexoff, J. Logan, C. Wong, K. Pradhan, E. C. Caparelli, Y. Ma, and M. Jayne, “Effects of low-field magnetic stimulation on brain glucose metabolism,” Neuroimage, vol. 51, no. 2, pp. 623–628, 2010.

[12] J. K. Deans, A. D. Powell, and J. G. Jefferys, “Sensitivity of coherent oscillations in rat hippocampus to AC electric fields,” J Physiol, vol. 583, no. Pt 2, pp. 555–565, 2007.

[13] J. T. Francis, B. J. Gluckman, and S. J. Schiff, “Sensitivity of neurons to weak electric fields,” J Neurosci, vol. 23, no. 19, pp. 7255–7261, 2003.

[14] W. A. Carlezon, Jr., M. L. Rohan, S. D. Mague, E. G. Meloni, A. Parsegian, K. Cayetano, H. C. Tomasiewicz, E. D. Rouse, B. M. Cohen, and P. F. Renshaw, “Antidepressant-like effects of cranial stimulation within a low-energy magnetic field in rats,” Biol Psychiatry, vol. 57, no. 6, pp. 571–576, 2005.

[15] M. Fava, M. P. Freeman, M. Flynn, B. B. Hoeppner, R. Shelton, D. V. Iosifescu, J. W. Murrough, D. Mischoulon, C. Cusin, M. Rapaport, B. W. Dunlop, M. H. Trivedi, M. Jha, G. Sanacora, G. Hermes, and G. I. Papakostas, “Double-blind, proof-of-concept (POC) trial of Low-Field Magnetic Stimulation (LFMS) augmentation of antidepressant therapy in treatment-resistant depression (TRD),” Brain Stimul, vol. 11, no. 1, pp. 75–84, 2018.

[16] M. L. Rohan, R. T. Yamamoto, C. T. Ravichandran, K. R. Cayetano, O. G. Morales, D. P. Olson, G. Vitaliano, S. M. Paul, and B. M. Cohen, “Rapid mood-elevating effects of low field magnetic stimulation in depression,” Biol Psychiatry, vol. 76, no. 3, pp. 186–193, 2014.

[17] M. Rohan, A. Parow, A. L. Stoll, C. Demopulos, S. Friedman, S. Dager, J. Hennen, B. M. Cohen, and P. F. Renshaw, “Low-field magnetic stimulation in bipolar depression using an MRI-based stimulator,” Am J Psychiatry, vol. 161, no. 1, pp. 93–98, 2004.

[18] G. M. Chandler, D. V. Iosifescu, M. H. Pollack, S. D. Targum, and M. Fava, “RESEARCH: Validation of the Massachusetts General Hospital Antidepressant Treatment History Questionnaire (ATRQ),” CNS Neurosci Ther, vol. 16, no. 5, pp. 322–325, 2010.

[19] M. Fava, “Diagnosis and definition of treatment-resistant depression,” Biol Psychiatr, vol. 53, no. 8, pp. 649–659, 2003.

[20] M. Fava and K. G. Davidson, “Definition and epidemiology of treatment-resistant depression,” Psychiatr Clin North Am, vol. 19, no. 2, pp. 179–200, 1996.

[21] D. H. Avery, P. E. Holtzheimer, 3rd, W. Fawaz, J. Russo, J. Neumaier, D. L. Dunner, D. R. Haynor, K. H. Claypoole, C. Wajdik, and P. Roy-Byrne, “A controlled study of repetitive transcranial magnetic stimulation in medication-resistant major depression,” Biol Psychiatry, vol. 59, no. 2, pp. 187–194, 2006.

[22] C. Loo, T. McFarquhar, and G. Walter, “Transcranial magnetic stimulation in adolescent depression,” Australas Psychiatry, vol. 14, no. 1, pp. 81–85, 2006.

[23] M. P. Fleck, M. F. Poirier-Littre, J. D. Guelfi, M. C. Bourdel, and H. Loo, “Factorial structure of the 17-item Hamilton Depression Rating Scale,” Acta Psychiatr Scand, vol. 92, no. 3, pp. 168–172, 1995.

[24] R. L. O’Sullivan, M. Fava, C. Agustin, L. Baer, and J. F. Rosenbaum, “Sensitivity of the six-item Hamilton Depression Rating Scale,” Acta Psychiatr Scand, vol. 95, no. 5, pp. 379–384, 1997.

[25] D. Watson, L. A. Clark, and A. Tellegen, “Development and validation of brief measures of positive and negative affect: the PANAS scales,” J Pers Soc Psychol, vol. 54, no. 6, pp. 1063–1070, 1988.

[26] R. A. Stern, VAMS : visual analog mood scales : professional manual. Odessa, FL: Psychological Assessment Resources, 1997.

[27] E. P. Ahearn, “The use of visual analog scales in mood disorders: a critical review,” J Psychiatr Res, vol. 31, no. 5, pp. 569–579, 1997.

[28] M. F. Folstein and R. Luria, “Reliability, validity, and clinical application of the Visual Analogue Mood Scale,” Psychol Med, vol. 3, no. 4, pp. 479–486, 1973.

[29] J. J. Daly, J. Prudic, D. P. Devanand, M. S. Nobler, S. H. Lisanby, S. Peyser, S. P. Roose, and H. A. Sackeim, “ECT in bipolar and unipolar depression: differences in speed of response,” Bipolar Disord, vol. 3, no. 2, pp. 95–104, 2001.

[30] A. U. Haq, A. F. Sitzmann, M. L. Goldman, D. F. Maixner, and B. J. Mickey, “Response of depression to electroconvulsive therapy: a meta-analysis of clinical predictors,” J Clin Psychiatry, vol. 76, no. 10, pp. 1374–1384, 2015.

[31] J. P. O’Reardon, H. B. Solvason, P. G. Janicak, S. Sampson, K. E. Isenberg, Z. Nahas, W. M. McDonald, D. Avery, P. B. Fitzgerald, C. Loo, M. A. Demitrack, M. S. George, and H. A. Sackeim, “Efficacy and safety of transcranial magnetic stimulation in the acute treatment of major depression: a multisite randomized controlled trial,” Biol Psychiatry, vol. 62, no. 11, pp. 1208–1216, 2007.

[32] M. Avissar, F. Powell, I. Ilieva, M. Respino, F. M. Gunning, C. Liston, and M. J. Dubin, “Functional connectivity of the left DLPFC to striatum predicts treatment response of depression to TMS,” Brain Stimul, vol. 10, no. 5, pp. 919–925, 2017.

[33] J. Downar, J. Geraci, T. V. Salomons, K. Dunlop, S. Wheeler, M. P. McAndrews, N. Bakker, D. M. Blumberger, Z. J. Daskalakis, S. H. Kennedy, A. J. Flint, and P. Giacobbe, “Anhedonia and reward-circuit connectivity distinguish nonresponders from responders to dorsomedial prefrontal repetitive transcranial magnetic stimulation in major depression,” Biol Psychiatry, vol. 76, no. 3, pp. 176–185, 2014.

[34] A. T. Drysdale, L. Grosenick, J. Downar, K. Dunlop, F. Mansouri, Y. Meng, R. N. Fetcho, B. Zebley, D. J. Oathes, A. Etkin, A. F. Schatzberg, K. Sudheimer, J. Keller, H. S. Mayberg, F. M. Gunning, G. S. Alexopoulos, M. D. Fox, A. Pascual-Leone, H. U. Voss, B. J. Casey, M. J. Dubin, and C. Liston, “Resting-state connectivity biomarkers define neurophysiological subtypes of depression,” Nat Med, vol. 23, no. 1, pp. 28–38, 2017.

[35] M. D. Fox, R. L. Buckner, M. P. White, M. D. Greicius, and A. Pascual-Leone, “Efficacy of transcranial magnetic stimulation targets for depression is related to intrinsic functional connectivity with the subgenual cingulate,” Biol Psychiatry, vol. 72, no. 7, pp. 595–603, 2012.

[36] C. Liston, A. C. Chen, B. D. Zebley, A. T. Drysdale, R. Gordon, B. Leuchter, H. U. Voss, B. J. Casey, A. Etkin, and M. J. Dubin, “Default mode network mechanisms of transcranial magnetic stimulation in depression,” Biol Psychiatry, vol. 76, no. 7, pp. 517–526, 2014.

[37] T. V. Salomons, K. Dunlop, S. H. Kennedy, A. Flint, J. Geraci, P. Giacobbe, and J. Downar, “Resting-state cortico-thalamic-striatal connectivity predicts response to dorsomedial prefrontal rTMS in major depressive disorder,” Neuropsychopharmacology, vol. 39, no. 2, pp. 488–498, 2014.

[38] A. Weigand, A. Horn, R. Caballero, D. Cooke, A. P. Stern, S. F. Taylor, D. Press, A. Pascual-Leone, and M. D. Fox, “Prospective validation that subgenual connectivity predicts antidepressant efficacy of transcranial magnetic stimulation sites,” Biol Psychiatry, vol. 84, no. 1, pp. 28–37, 2018.

[39] Z.-D. Deng, S. H. Lisanby, and A. V. Peterchev, “Electric field depth–focality tradeoff in transcranial magnetic stimulation: simulation comparison of 50 coil designs,” Brain Stimul, vol. 6, no. 1, pp. 1–13, 2013.

[40] B. Wang, M. R. Shen, Z.-D. Deng, J. E. Smith, J. J. Tharayil, C. J. Gurrey, L. J. Gomez, and A. V. Peterchev, “Redesigning existing transcranial magnetic stimulation coils to reduce energy: application to low field magnetic stimulation,” J Neural Eng, vol. 15, no. 3, 036022, 2018.

